# SASH1 impairs melanin synthesis and metastasis by down-regulating the TGF-β signaling pathway

**DOI:** 10.1101/2024.05.22.595382

**Authors:** Hongzhou Cui, Qiong Wang, Honggang Liang, Yingjie Zhang, Bo Liang, Wenjun Wang, Shanshan Ge, Hongxia He, Xiaoli Ren, Zhenxing Su, Shuping Guo

**Author notes:** **Corresponding author,** Shuping Guo, Department of Dermatology, the First Hospital, Shanxi Medical University, Taiyuan, Shanxi, China. These authors contributed equally to this work (Hongzhou Cui, Qiong Wang, Honggang Liang).

## Abstract

Dyschromatosis universalis hereditaria (DUH) is a rare genetic dermatosis characterized by widespread hyperpigmentation and depigmentation. In our previous study, we identified SH3 domain-containing protein 1 (SASH1) mutations associated with the DUH phenotype in Chinese families and predict SASH1/*THBS1/*TGF-β1 signaling to mediate melanin production and melanocyte transport. We detected SASH1/*THBS1/*TGF-β1 pathway genes expression and evaluated cell phenotypes and melanin synthesis in A375 and PIG1 cell lines by regulating the expression of SASH1 and THBS1 respectively. Finally, we validated the results of the cellular experiments in nude mice. Results showed that SASH1 inhibited the proliferation, migration, invasion, EMT ability and melanin synthesis via TGF-β1 signaling, and THBS1 reversed the elevation of TGF-β1 caused by SASH1 knockdown. We demonstrated that SASH1 further inhibits TGF-β1 through its regulatory effect on THBS1, thereby inhibiting melanin synthesis and metastasis, which may advance the utilization of TGF-β1 for therapeutic purposes.

**Plain Language Summary:** Dyschromatosis universalis hereditaria(DUH) is a genetic disease showing uneven pigment pattern, which greatly affects the appearance. In our previous study, we had reported a SASH1 mutation in a Chinese DUH pedigree. In this study, we mainly focus on the effect of SASH1 gene on pigment synthesis and metabolism at the cell and animal level. The results show SASH1 inhibits the melanin synthesis of melanocytes, it can also hinder cell migration, invasion, and EMT. More than that, SASH1 reduced TGF-β1 mRNA, protein expression and promoter activity, and THBS1 could discharge this effect. In mice, SASH1 inhibits the tumor growth via TGF-β1 signaling. We detected a novel SASH1/*THBS1/*TGF-β1 pathway in cell phenotypes and melanin synthesis.

## INTRODUCTION

Dyschromatosis universalis hereditaria (DUH) is a rare genetic dermatosis characterized by widespread hyperpigmentation and depigmentation.(Cao et al. 2021) The typical lesions most often occur within the first year of life, and the pigmentation abnormalities appear first on the trunk or extremities, spreading to all parts of the body, including the face, neck, palms, and plantar areas, and mucous membranes over the few years.(Sethuraman et al. 2002; Wu et al. 2020) In some cases, nail, dental, and hair involvement may also be observed.(Al Hawsawi et al. 2002; Bukhari et al. 2006) The lesions usually stop expanding after puberty but do not fade independently.(Wu et al. 2020) Melanocytes are a minority cell population within the basilar epidermis, primarily providing melanin pigment to their neighboring keratinocytes. When melanin production is compromised, it leads to hypopigmentation or hyperpigmentation. Previous studies have shown that the number of melanocytes is not affected in patients with autosomal recessive DUH, unlike the synthesis and maturation of melanosomes in melanocytes.(Gupta et al. 2015) Treatment of patients with DUH remains limited as the effects on the biological behavior of melanocytes are still poorly understood.(Zhang et al. 2017) Therefore, early detection of their pathogenic mechanisms is essential for improving the diagnosis and treatment, and a better understanding of the molecular mechanisms of melanocyte genesis and development in DUH may help to explore new sensitive and specific biomarkers and therapeutic drugs.

In recent years, significant progress has also been made in genetic-related research. Gene detection techniques have shown that mutations in the *ABCB6* gene and *SASH1* gene are closely related to DUH.(Cao et al. 2021; Liu et al. 2021; Lu et al. 2014; Wu et al. 2020; Zhong et al. 2019) In our previous study, we identified SASH1 mutations associated with the DUH phenotype in Chinese families and used synthetic modeling of mRNA expression profiles to predict that SASH1/TGF-β1 signaling is involved in cell migration and that *THBS1* may be involved in this cascade to regulate melanin production and melanocyte transport.(Cui et al. 2020) Chen *et al*. found that downregulation of the *SASH1* gene results in higher migration and invasion of A549 cells.(Chen et al. 2012) Zhou *et al*. showed that *SASH1* variants may disrupt melanocyte-keratinocyte adhesion by decreasing E-calmodulin expression, increasing A375 cell motility and altering melanin transfer.(Zhou et al. 2013) The transforming growth factor β (TGF-β) signaling pathway has an important role in the epithelial-to-mesenchymal transition (EMT), and EMT regulates tumor morphogenesis and progression.(Derynck et al. 2021) However, the underlying upstream mechanism of TGF-β signaling in the generation of melanocytes remains largely unclear. THBS1 was found to be significantly upregulated in melanocyte samples from NBUVB-treated augmentations, suggesting that it is involved in cell adhesion and maintenance of the stemness of melanocyte precursors in the augmentations via β-catenin signaling.(Goldstein et al. 2018) However, the roles of *SASH1* overexpressed THBS1 and suppressed TGF-β1 signaling in melanocytes are rarely studied.

This study explored the role of SASH1-dependent regulating melanin synthesis and metastasis using both *in vitro* and *in vivo* approaches. qRT-PCR, Western blot, ELISA, and CCK8 were used for the experiments. We found that SASH1 impairs melanin synthesis and metastasis by affecting the TGF-β signaling pathway through THBS1.

## MATERIALS AND METHODS

### Cell culture

Human melanoma cell line A375 and normal melanocyte PIG1 were obtained from the American Type Culture Collection (ATCC, Manassas, VA, USA). These cells were maintained in a DMEM (Thermo Fisher Scientific, Waltham, MA, USA), supplemented with 10% FBS (Gibco, USA) and 1% penicillin-streptomycin (Gibco, USA). Cells were cultured in a humidified incubator at 37°C with 5% CO2 (Sanyo Osaka, Japan).

### Establishment of stable SASH1/THBS1 knock-down and overexpression A375/PIG1 cell lines

To construct stable SASH1/THBS1 knock-down A375/PIG1 cell lines, shRNAs specific targeting to SASH1/THBS1 was designed by Invitrogen online tool (https://rnaidesigner.thermofisher.com/rnaiexpress/) and cloned into the lentiviral pLVX-sh1 vector. Lentivirus carrying SASH1/THBS1 shRNAs were constructed by Genelily Biotech Co., LTD (Shanghai, China). LV-NC represents an empty vector packaged by lentivirus as negative controls. The A375/PIG1 cell lines in the 6-well plate were infected with the lentivirus following the manufacturer’s instruction. After 24 hours, the medium was replaced with a complete medium. The stably infected A375/PIG1 cells were selected by incubation with 2 μg/ml of puromycin for two weeks. The efficiency of SASH1/THBS1 knock-down was examined by qRT-PCR and Western blot.

In order to construct stable SASH1/THBS1 overexpression A375/PIG1 cell lines, the human SASH1/THBS1 coding sequences were purchased from Generalbiol Biotech (Hefei, Anhui, China) and cloned into the lentiviral pLVX-IRES-ZsGreen vector. Lentivirus carrying SASH1/THBS1 coding sequences were constructed by Genelily Biotech Co., LTD (Shanghai, China). The vector control or SASH1 knock-down cell lines were seeded in a 6-well plate and infected with the lentivirus following the manufacturer’s instruction. After 24 hours, the medium was replaced by using the complete medium. The stably infected ZsGreen A375/PIG1 cells were isolated by fluorescence-activated cell sorting (FACS), after which the efficiency of SASH1/THBS1 overexpression was examined by qRT-PCR and Western blot.

### Real-time PCR

Total RNA was extracted using TRIzol reagent (Invitrogen, Carlsbad, CA, USA) and reverse-transcribed using the cDNA Synthesis kit (Takara, Japan). The RNA quantity and density were verified by a spectrophotometer. qRT-PCR was performed using the SYBR Green Master MIX Kit (Takara, Japan) according to the manufacturer’s instructions. The assays were run in triplicate, and relative gene expression was determined by using the 2^-ΔΔCt^ method.

### Western Blot

For total protein extraction, cell lysates were obtained using RIPA buffer (Beyotime, Shanghai, China) supplemented with phosphatase and protease inhibitors (Yeasen, Shanghai, China). A total of 15 μL protein was injected into a Bis-Tris SDS/PAGE gel and transferred to PVDF membranes. After blocking with 5% BSA, the membranes were incubated overnight at 4°C with primary antibody: anti-SASH1 antibody (Abcam, UK, 1:1000), anti-E-cadherin antibody (Abcam, UK, 1:2000), anti-Vimentin antibody (Abcam, UK, 1:2000), anti-TGF-β antibody (Abcam, UK, 1:2000), anti-GAPDH antibody (Boster, China, 1:5000), after which they were exposed to HRP Conjugated AffiniPure Mouse Anti-human IgG (H+L) (Boster, China, 1:5000) for 60 min. Bands were incubated with an ECL kit and analyzed with an imaging system. Densitometric analysis was carried out using Adobe Photoshop CS6.

### Enzyme-linked immunosorbent assay (ELISA)

ELISA for cytokines detection was performed with kits (Boster Bio, Wuhan, China).

### CCK-8 assay

A total of 5× 10^3^ cells were inoculated into each well of 96-well plates. At each time point (24, 48, 72, and 96 h), 10 μL sterile CCK-8 solution (Yeasen, Shanghai, China) was added to the sextuplicate wells. The wells were incubated for 3 h, and the absorbance of each well was determined at 450 nm.

### Wound-healing migration assay

The cells were cultured in 6-well plates at 5×10^5^ per well. Cell monolayers were mechanically disrupted using a sterile 200 μl micropipette tip to generate a linear wound. The average distance migrated by the cells was measured using a microscope (Nikon, ECLIPSE Ts2) calibrated with an ocular micrometer.

### Invasion assay

Cells were incubated using 24-well transwell plates (8 μm pore size, Corning, NY, USA). One million cells suspended in a serum-free medium were plated in the upper chambers with Matrigel (BD Biosciences, USA), and a 0.6 ml medium with 10% FBS was added to the lower chamber. After incubation for a suitable amount of time, the cells were fixed in 4% paraformaldehyde, stained with crystal violet, and counted under a microscope.

### Melanin determination

Cells were cultured at 37℃, 5% CO_2_ for 24 h. After 48 hours, they were washed with PBS, harvested with trypsin, centrifuged at 1500 × g for 10 min, dissolved in 1 M NaOH containing 10% DMSO, and incubated at 80℃ for 2 h. Melanin content was determined by spectrophotometry at 475 nm with Microplate Reader (Molecular Devices, USA) and expressed as relative absorbance unit /10^5^ cells.

### In vivo assays

All animal experiments were approved by the Animal Experimentation Ethics Committee of the first affiliated Hospital, Shanxi Medical University. Mice were housed at 22 °C on a 12-hour light/dark cycle in individual cages and fed ad libitum with standard food and deionized water. Six-week-old C57BL/6 mice were maintained according to the guidelines of the 3Rs (replacement, reduction, and refinement).

A total of 5 × 10^6^ cells resuspended in 100 μL of PBS were inoculated subcutaneously in the left flank of the mice. Mice were divided into four groups (5 mice per group): shNC group, shSASH1 group, shNC+anti-TGF-b1 group, and shSASH1+anti-TGF-b1 group. After the tumor was detected, tumor size was measured every 3 days by a vernier caliper, and tumor volume was calculated as volume (cm3) = L x W^2^ × 0.5, with L and W representing the largest and smallest diameters, respectively.

### Statistical analysis

Statistical analyses were performed using the SPSS software version 19.0 (SPSS, Inc., Chicago, IL) or GraphPad Prism 7.0 (GraphPad Software, USA) as previously described. The values are presented as the mean ± standard deviation (SD). The chi-squared test, student’s t-test, Mann-Whitney U test, Kruskal-Wallis test, and one-way ANOVA test were used for group comparisons, as appropriate. The survival curve was prepared using the Kaplan-Meier method and analyzed by the log-rank t-test. Cox’s proportional hazard regression model was used to analyze the independent prognostic factors. P < 0.05 represented statistical significance.

## RESULTS

### SASH1 inhibits the proliferation ability in melanocytes

In order to evaluate the functional role of SASH1 in melanoma cells, we first knock-downed the expression of SASH1 in melanoma cell A375 and immortalized human melanocyte cells (PIG1) via lentivirus. As shown in **Figure 1A** (all p<0.05), the mRNA level of SASH1 was dramatically decreased by SASH1 shRNA expressing lentivirus. The protein levels of SASH1 were further confirmed by Western Blot (all p<0.05, **Figure 1B**). Furthermore, the CCK-8 assay showed that both A375 and PIG1 cells’ proliferation ability improved after SASH1 knock-down (**Figure 1C**).

**Figure 1.**
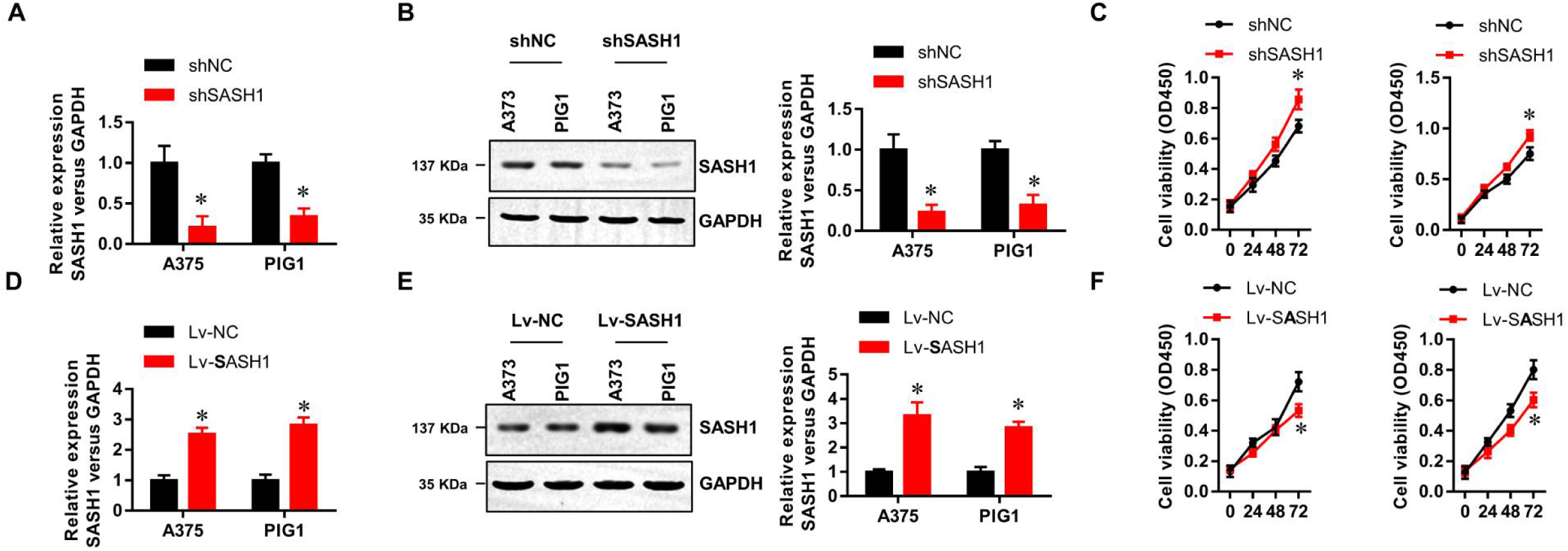
SASH1 inhibits the proliferation ability in melanocytes. **(A, D)** The mRNA level of SASH1 in A375 and PIG1 cells was detected by real-time PCR. (**B, E)** The protein levels of SASH1 in A375 and PIG1 cells detected by Western Blot. **(C, F)** The cell viability of A375 and PIG1 cells analyzed by CCK-8. *P<0.05 vs. negative control (NC) group.

Next, we overexpressed the SASH1 in A375 and PIG1 cells via the SASH1 coding sequence expressing lentivirus. The mRNA (p<0.05, **Figure 1D**) and protein (p<0.05, **Figure 1E**) levels were evaluated by real-time PCR and Western Blot. Consistently, the proliferation ability of both A375 and PIG1 cells was impaired by SASH1 overexpression (**Figure 1F**).

### SASH1 inhibits the migration, invasion, and EMT in melanoma cells

We performed a wound healing assay and Transwells invasion assay to further analyze the effect of SASH1 on metastasis in melanoma cells. As shown in **Figure 2A**, SASH1 knock-down significantly enhanced the migration ability in A375 cells, whereas SASH1 overexpression inhibited the migration ability in A375 cells (all p<0.05). Similarly, SASH1 knock-down significantly elevated the invasion ability in A375 cells, which was then inhibited by SASH1 overexpression (all p<0.05, **Figure 2B**). Moreover, the epithelial-to-mesenchymal transition (EMT) was activated by SASH1 knock-down, indicated by the decreased level of E-cadherin and increased level of vimentin (both p<0.05, **Figure 2C**). These results demonstrate that SASH1 inhibits melanoma cells’ migration, invasion, and EMT.

**Figure 2.**
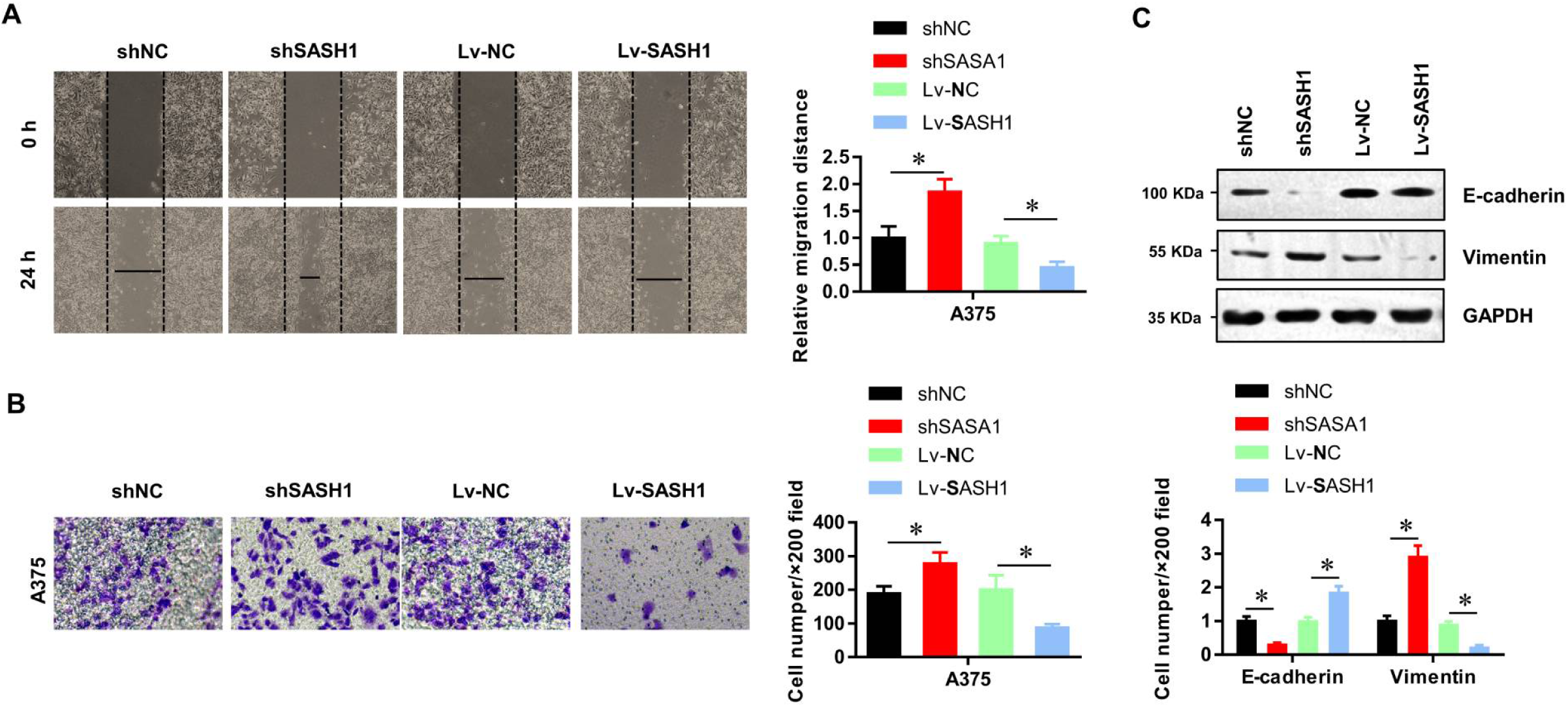
SASH1 inhibits the migration, invasion, and EMT in melanoma cells. **(A)** The migration ability of A375 and PIG1 cells was detected by wound healing assay. **(B)** The invasion ability of A375 and PIG1 cells detected by transwell assay. **(C)** The expression markers of EMT analyzed by Western Blotting; *P<0.05 vs. negative control (NC) group.

### SASH1 inhibits the melanin synthesis of melanocytes

In order to further explore the effects of SASH1 on melanin synthesis of melanocytes, melanin content assays were performed. As shown in **Figure 3** (all p<0.05), SASH1 knock-down significantly decreased the melanin content in PIG1 cells, whereas SASH1 overexpression inhibited the melanin content in PIG1 cells.

**Figure 3.**
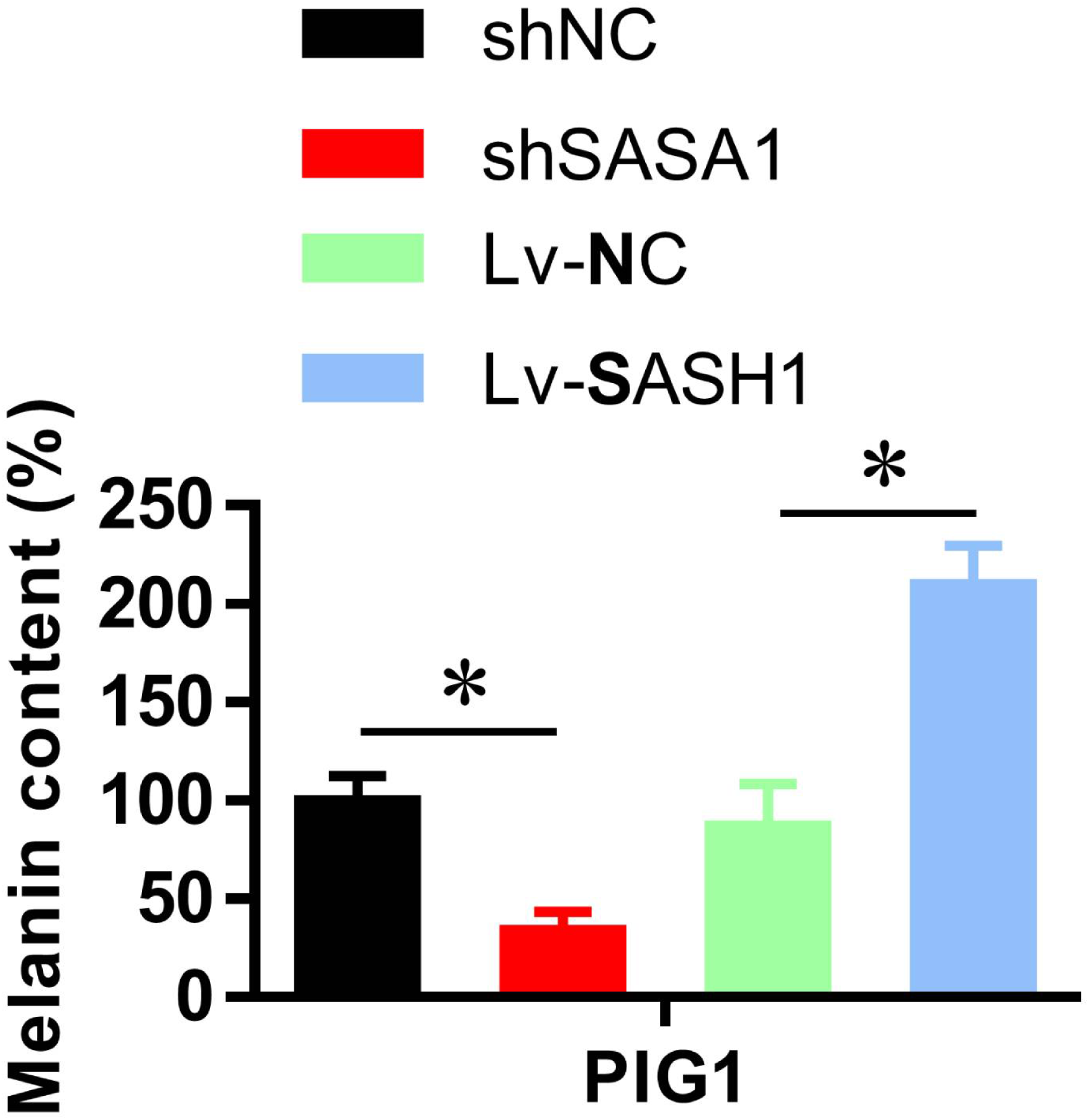
SASH1 inhibits the melanin synthesis of melanocytes. *P<0.05 vs. negative control (NC) group.

### SASH1 transcriptionally inhibits the level of TGF-β1

We previously reported that SASH1 mutation inhibits the TGF-β1 signaling in PIG1 cells. Subsequently, we tried to discover whether SASH1 regulates the TGF-β1 signaling in melanoma cells. As shown in **Figure 4A**, the mRNA level of TGF-β1 in SASH1 knock-downed A375 and PIG1 cells was consistently enhanced (A375: p<0.01, PIG1: p<0.05), whereas it was decreased by SASH1 over-expression (A375: p<0.001, PIG1: p<0.01, **Figure 4B**). Similarly, the secreted TGF-β1 protein was elevated by SASH1 knock-down (**Figure 4C**) and decreased by SASH1 overexpression (**Figure 4D**), which was further confirmed by Western blot (**Figure 4E**).

**Figure 4.**
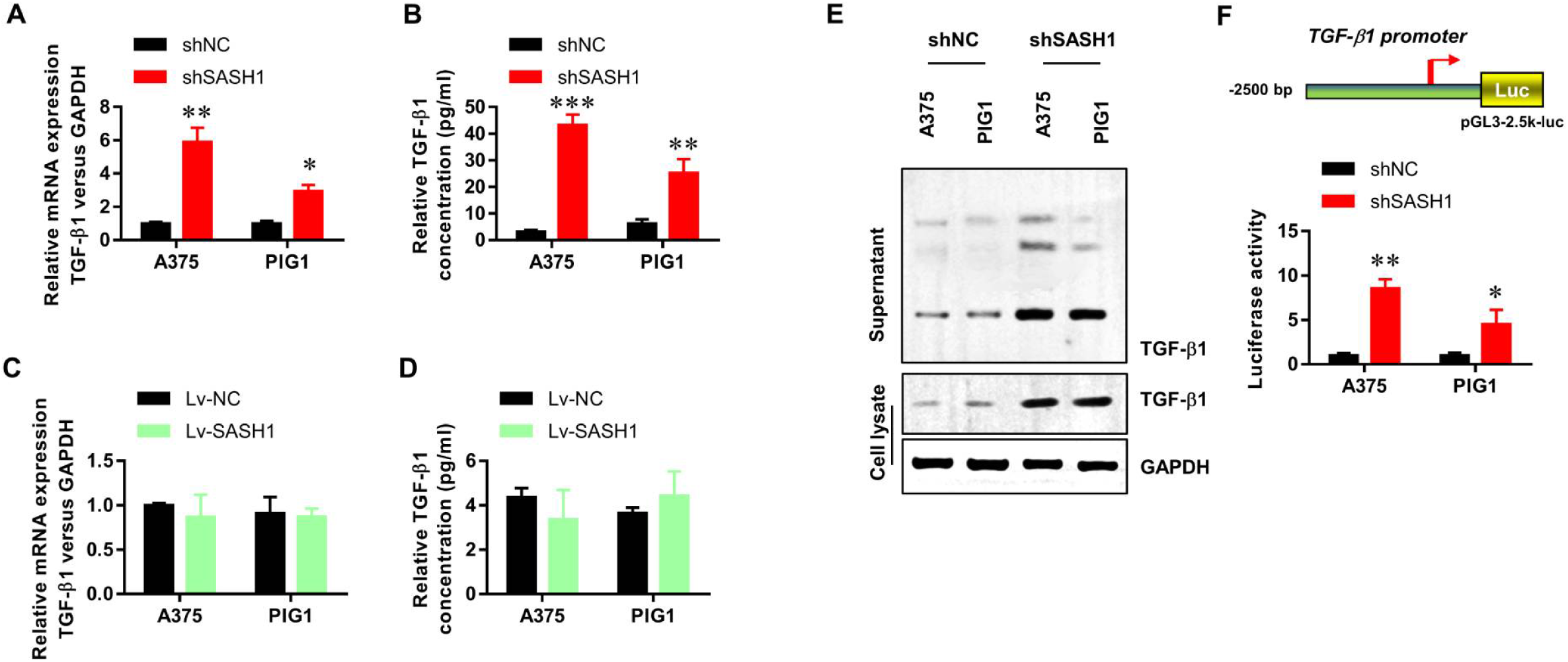
SASH1 transcriptionally inhibits the level of TGF-β1. **(A, C)** The mRNA level of TGF-β1 in A375 and PIG1 cells detected by real-time PCR. **(B, D)** The protein levels of secreted TGF-β1 in A375 and PIG1 cells detected by ELISA. **(E)** The protein levels of secreted TGF-β1 in A375 and PIG1 cells were confirmed by Western Blot. **(F)** The promoter activity of TGF-β1 detected by the dual-luciferase assay. *P<0.05 vs. negative control (NC) group, **P<0.01 vs. negative control (NC) group, ***P<0.001 vs. negative control (NC) group.

Next, we performed a dual-luciferase reporter assay to evaluate the promoter activity of TGF-β1 in these A375 and PIG1 cells. The results showed that the promoter activity of TGF-β1 was also elevated by SASH1 knock-down and decreased by SASH1 overexpression (A375: p<0.01, PIG1: p<0.05, **Figure 4F**). These results suggest that SASH1 transcriptionally inhibits the level of TGF-β1.

### THBS1 reversed the elevation of TGF-β1 caused by SASH1 knockdown

The release of biologically active TGF-β1 isoform from a latent complex involves proteolytic processing of the complex and/or induction of conformational changes by proteins such as thrombospondin-1 (THBS1). We overexpressed THBS1 in A375 and PIG1 cells with or without SASH1 knock-down. As shown in **Figure 5A**, THBS1 overexpression did not affect the expression level of TGF-β1 mRNA in control A375 and PIG1 cells (all p>0.05), whereas overexpression of THBS1 reversed the decrease in TGF-β mRNA level due to knock-down of SASH1, suggesting that SASH1 reduces the expression level of TGF-β through its regulatory effect on THBS1 (all p<0.05). ELISA (**Figure 5B**) and Western Blotting (**Figure 5C**) further confirmed a similar observation in the protein level of TGF-β1. Furthermore, the promoter activity of TGF-β1 was similarly inhibited by THBS1 in the SASH1 knock-down A375 and PIG1 cells (**Figure 5D**).

**Figure 5.**
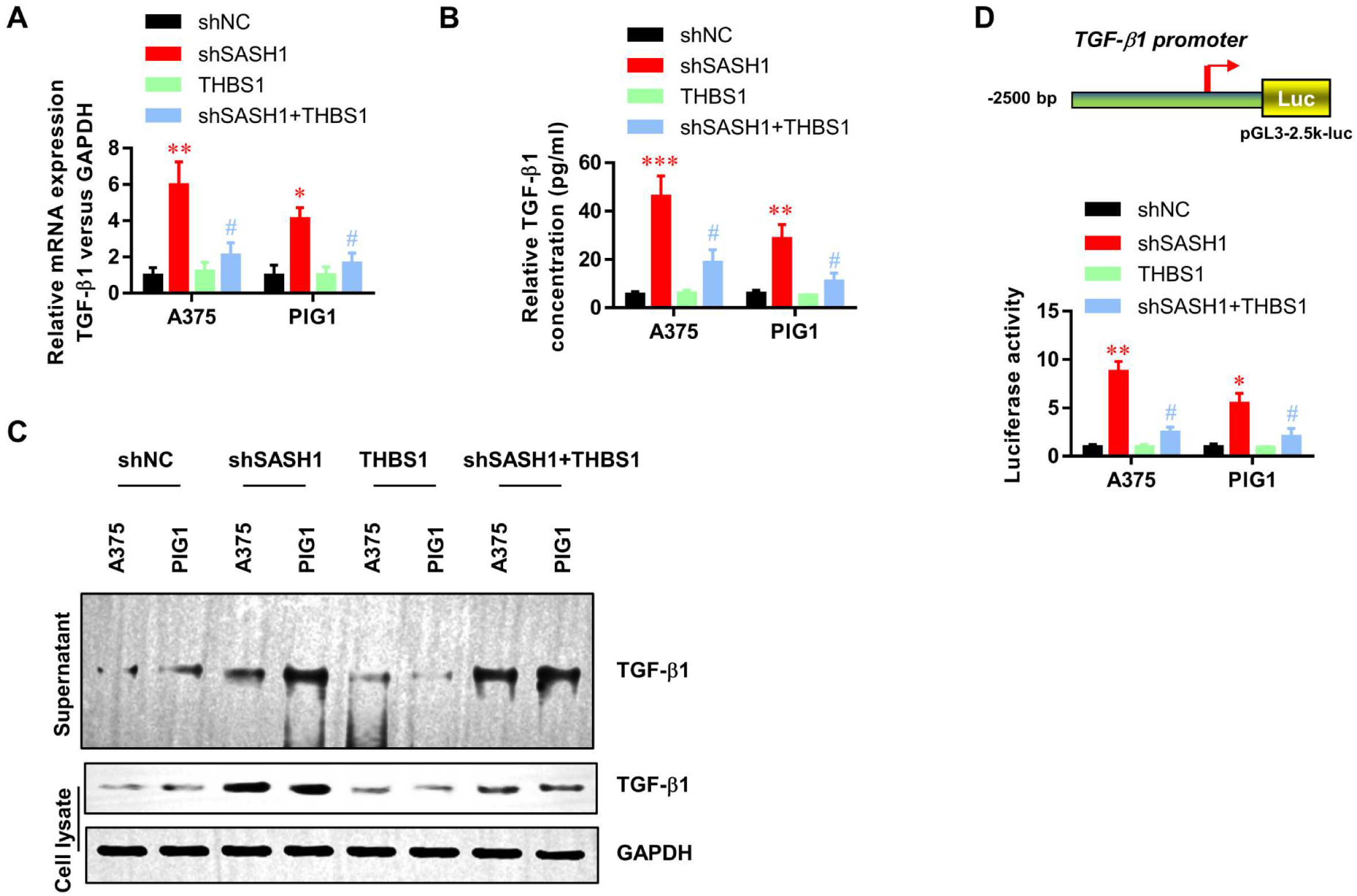
THBS1 reversed the SASH1 knock-down elevated expression of TGF-β1. **(A)** The mRNA levels of TGF-β1 in A375 and PIG1 cells detected by real-time PCR. **(B)** The protein levels of secreted TGF-β1 in A375 and PIG1 cells detected by ELISA. **(C)** The protein levels of secreted TGF-β1 in A375 and PIG1 cells confirmed by Western Blotting. (D) The promoter activity of TGF-β1 detected by dual-luciferase assay. *P<0.05 vs. negative control (NC) group, ^#^P<0.05 vs. SASH1 group.

### SASH1 inhibits the proliferation, migration, and invasion abilities via TGF-β1 signaling in melanoma cells

To verify whether SASH1 regulates the biological function of melanoma cells through TGF-β1 signaling, we neutralized SASH1 knock-down elevated TGF-β1 with antibodies in A375 and PIG1 cells. The anti-TGF-β1 antibody did not affect the proliferation (**Figure 6A**), migration (**Figure 6B**), and invasion (**Figure 6C**) of normal A375 and PIG1 cells (all p>0.05). On the contrary, we added recombinant TGF-β1 protein in A375 and PIG1 cells with or without SASH1 overexpression, finding that TGF-β1 dramatically enhanced proliferation (p<0.05, **Figure 6A**), migration (p<0.01, **Figure 6B**), and invasion (p<0.01, **Figure 6C**) in A375 and PIG1 cells. These results suggest that SASH1 inhibits the proliferation, migration, and invasion abilities via TGF-β1 signaling in melanoma cells.

**Figure 6.**
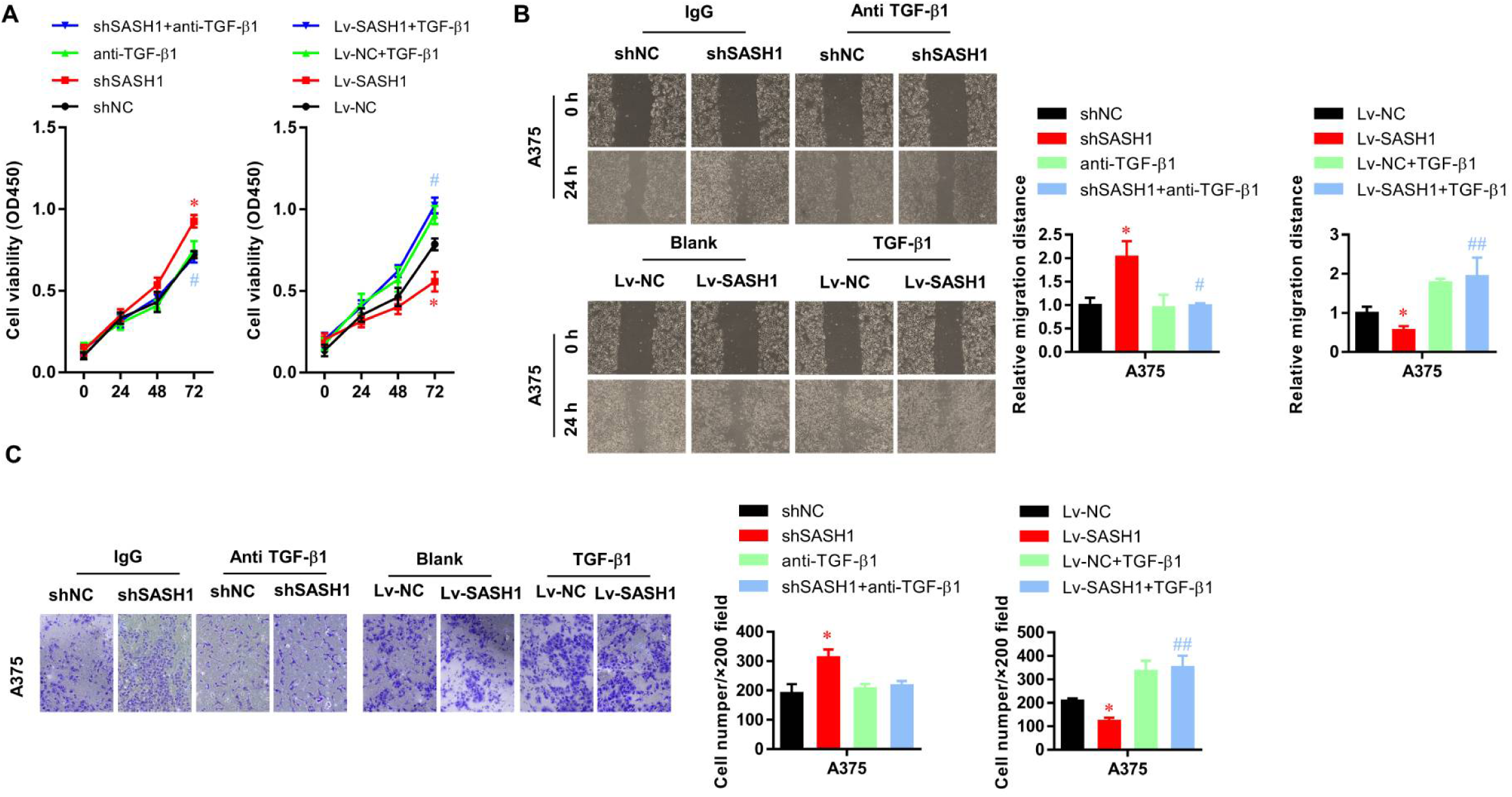
SASH1 inhibits the proliferation, migration, and invasion abilities via TGF-β1 signaling in melanoma cells. **(A)** The cell viability of A375 and PIG1 cells analyzed by CCK-8 assay. **(B)** The migration ability of A375 and PIG1 cells detected by wound healing assay. **(C)** The invasion ability of A375 and PIG1 cells detected by transwell assay. *P<0.05 vs. negative control (NC) group, ^#^P<0.05 vs. SASH1 group, ^##^P<0.01 vs. SASH1 group.

### SASH1 inhibits the melanin synthesis of melanocytes via TGF-β1 signaling

In order to further identify whether SASH1 regulates the melanin synthesis of melanoma cells via TGF-β1 signaling, melanin content assays were performed. As shown in **Figure 7**, the anti-TGF-β1 antibody did not affect the melanin content (p>0.05, **Figure 7A**) in normal PIG1 cells, whereas it reversed the SASH1 knock-down decreased melanin content (all p<0.05). Similarly, TGF-β1 did not affect the melanin content, while it reversed the SASH1 overexpression of elevated melanin content (**Figure 7B**).

**Figure 7.**
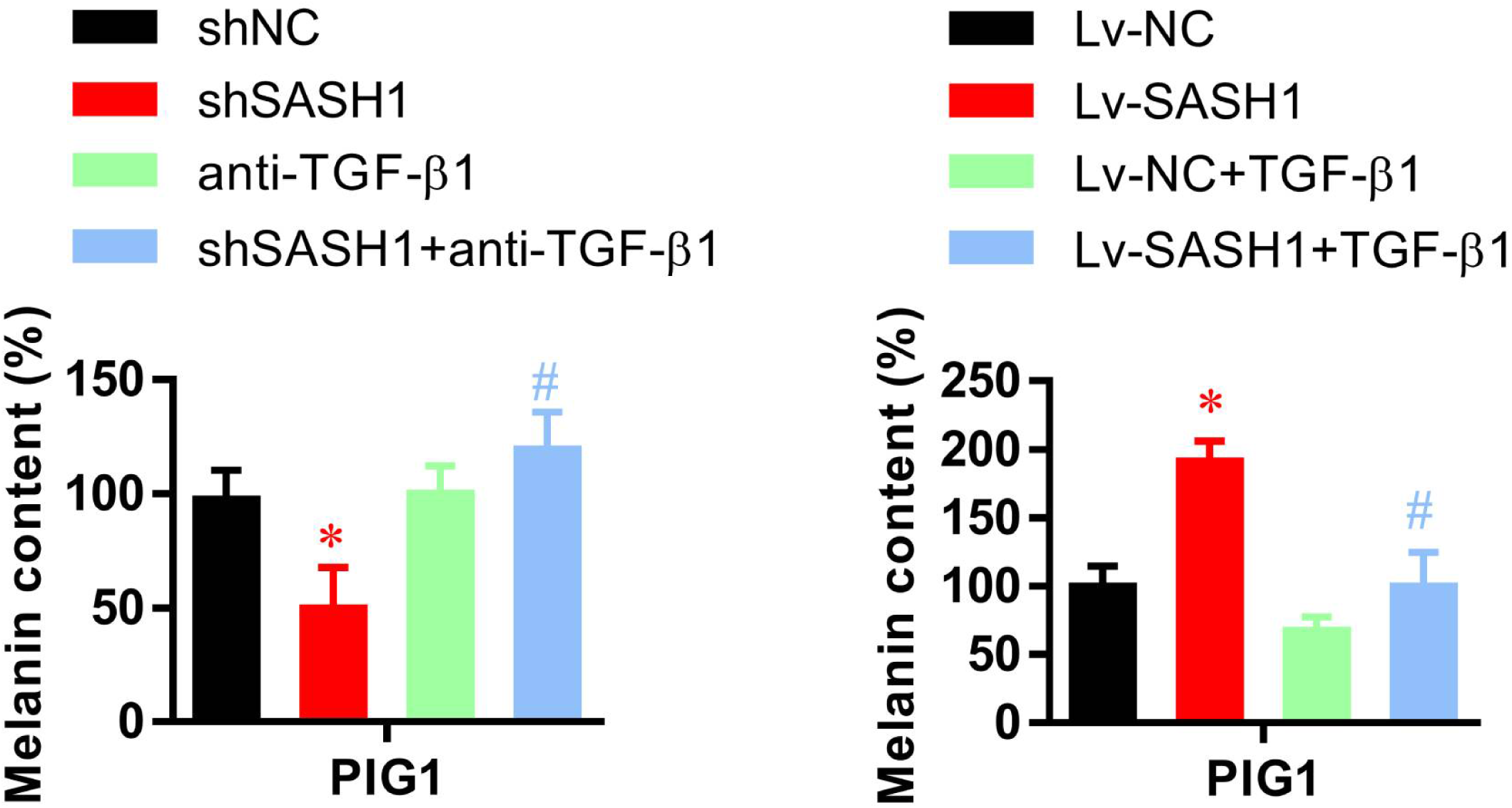
SASH1 inhibits the melanin synthesis of melanocytes via TGF-β1 signaling. *P<0.05 vs. negative control (NC) group, ^#^P<0.05 vs. SASH1 group.

### SASH1 inhibits the tumor growth via TGF-β1 signaling *in vivo*

To verify the biological effect of SASH1-TGF-β1 signaling *in vivo*, Balb/c NOD mice were subcutaneously implanted with A375 melanoma cells with or without SASH1 knock-down, and were divided into four groups (**Figure 8A**). Similar to the *in vitro* experiment, SASH1 knock-down significantly improved tumor growth (**Figure 8B**) and tumor weight (p<0.05, **Figure 8C**). Interestingly, anti-TGF-β1 antibody treatment dramatically inhibited tumor growth (**Figure 8B**) and decreased tumor weight (p<0.05, **Figure 8C**), which was different from the results observed in the *in vitro* study. Moreover, Anti-TGF-β1 antibody treatment also reversed the SASH1 knock-down and improved tumor growth (**Figure 8B**) and tumor weight (p<0.05, **Figure 8C**). Collectively, these results demonstrate that SASH1 inhibits tumor growth via TGF-β1 signaling *in vivo*.

up.

**Figure 8.**
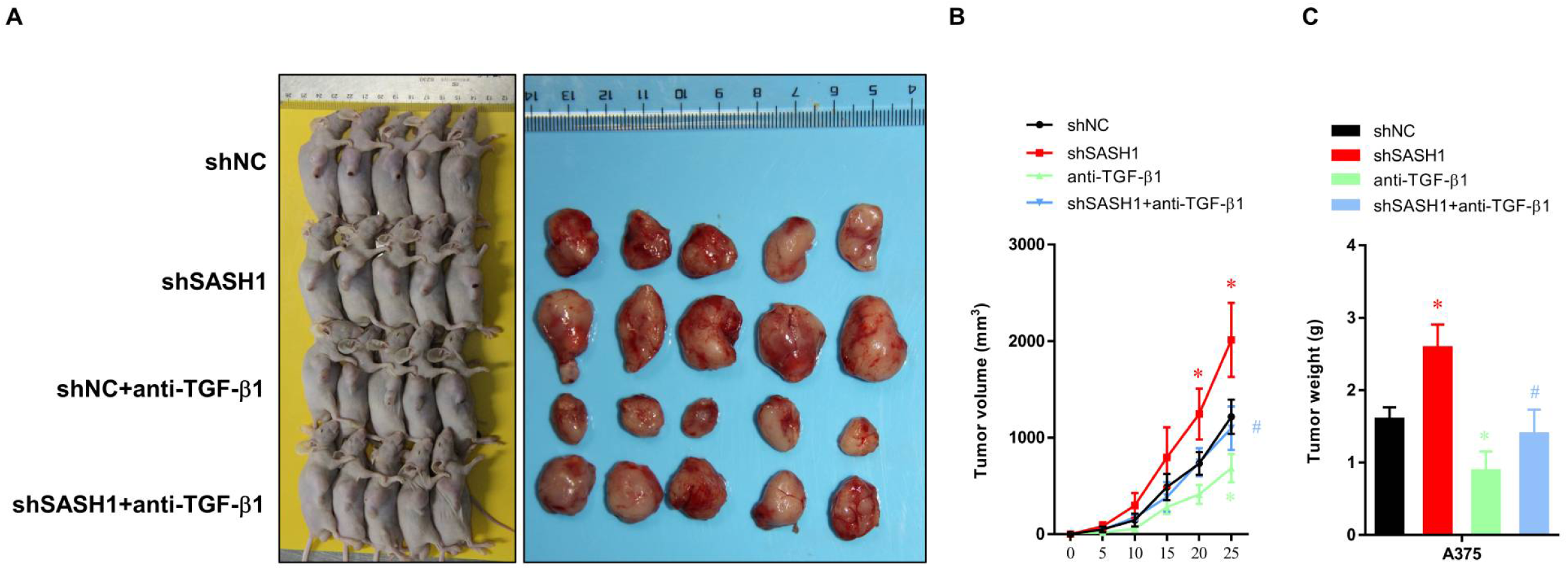
SASH1 inhibits tumor growth via TGF-β1 signaling *in vivo.* **(A)** The image of xenograft mice and tumors. **(B)** The tumor volume and **(C)** tumor weight in these four groups. *P<0.05 versus the negative control (NC) group, ^#^P<0.05 vs. SASH1 group.

## DISCUSSION

As a potential tumor suppressor gene, *SASH1* is reduced in various diseases.(Burgess et al. 2020; Gong et al. 2017; Lin et al. 2012) It has been reported that SASH1 has a crucial role in tumor occurrence, development, invasion, and metastasis.(Burgess et al. 2016) According to the characteristics of the SASH1 protein domain, SASH1 may have an important role in intracellular signal transduction and transcriptional regulation. The decrease in SASH1 expression is associated with the formation of heterogeneous distant metastasis. Another study demonstrated that SASH1 interacts with the actin cytoskeleton to stimulate cell-matrix adhesion.(Xie et al. 2017; Zhu et al. 2019) In melanoma, SASH1 derives a G2/M arrest in A375 cells.(Lin et al. 2012) However, there are limited studies on the role of SASH1 in normal melanocyte development. Therefore, we investigated the role of SASH1 in melanocytes *in vitro*. In this study, we demonstrated that SASH1 inhibited the proliferation, migration, and invasive ability of melanoma cells *in vitro* and tumor growth *in vivo* but also significantly inhibited the ability of melanin synthesis and metastasis in normal human skin melanocytes.

TGF-β1 is a multifunctional growth factor that has a key role in the development and tissue homeostasis by regulating cell proliferation, differentiation, and apoptosis in various cells.(Gumienny and Padgett 2002; Lutz and Knaus 2002; Syed 2016) However, little attention has been paid to the mechanisms of TGF-β-induced hypopigmentation. It has been suggested that TGF-β1 inhibits melanin synthesis by delaying ERK activation and subsequently reducing microphthalmia-associated transcription factor and tyrosinase production.(Kim et al. 2004) Xing *et al*. suggested that tranexamic acid stimulates TGF-β1 expression in keratinocytes, further inhibiting melanogenesis through paracrine signaling.(Xing et al. 2022) Herein, we found that SASH1 inhibits the proliferation, migration, and invasion ability of melanoma cells and melanocytes in vitro and tumor growth in vivo via TGF-β1 signaling. With the neutralizing of TGF-β1 by anti-TGF-β1 antibody, the tumor enhancer role and the inhibition of melanogenic capacity of SASH1 were reversed.

THBS1 is a multifunctional stromal ECM and secretory protein that is rich in platelet β granules and widely upregulated in tissue injury and repair.(Murphy-Ullrich and Suto 2018) It has a variety of specific receptor/binding partners and cellular functional domains.(Anastasi et al. 2020) THBS1 is a major regulator of TGF-β activation, which also has TGF-β-independent functions in hemostasis, cell adhesion, migration, and regulation of growth factors (EGF, VEGF, and FGF), inhibition of angiogenesis, and nitric oxide signal transduction.(Ouyang et al. 2020; Pal et al. 2016) Herein, we found that THBS1 overexpression did not affect the expression level of TGF-β1 mRNA and protein in normal A375 and PIG1 cells. In contrast, it dramatically abrogated the SASH1 knock-down elevated TGF-β1 expression.

In conclusion, we identified a novel mechanism through which SASH1 regulates melanin production. Gene mutation SASH1 caused downregulation of SASH1, leading to elevated expression of TGF-β1, which in turn promoted cell migration, cell invasion, EMT, and inhibited melanogenic capacity and the occurrence of DUH. Moreover, THBS1 overexpression was found to significantly attenuate the whole process. We elucidated the mechanism of the abnormal pigmentation of DUH caused by SASH1, which may advance the utilization of TGF-β1 for treatment.

## Statements and Declarations

### Author contributions

Conceptualization:SG

Data Curation: BL, HL and ZS

Formal Analysis: HH and XR

Funding Acquisition: SG

Methodology: WW, and YZ

Writing - Original Draft Preparation: HC and QW

Writing - Review and Editing: All authors.

### Data Availability Statement

All data generated or analysed during this study are included in this published article.

### Ethics approval

The authors are accountable for all aspects of the work in ensuring that questions related to the accuracy or integrity of any part of the work are appropriately investigated and resolved. Experiments were performed in compliance with the Animal [Scientific Procedures] Act 1986, national or institutional guidelines for the care and use of animals. The study adhered to all institutional and national guidelines for the care and use of laboratory animals. All animal experiments were approved by the Ethics Committee of Shanxi Medical University (2015LL045).

### Consent to participate

Not applicable.

### Consent to publish

Not applicable.

## Acknowledgements

None.

## References

Al Hawsawi K, Al Aboud K, Ramesh V, Al Aboud D. 2002. Dyschromatosis universalis hereditaria: Report of a case and review of the literature. Pediatric dermatology. 19(6):523–526.

Anastasi C, Rousselle P, Talantikite M, Tessier A, Cluzel C, Bachmann A, Mariano N, Dussoyer M, Alcaraz LB, Fortin L et al. 2020. Bmp-1 disrupts cell adhesion and enhances tgf-β activation through cleavage of the matricellular protein thrombospondin-1. Science signaling. 13(639).

Bukhari IA, El-Harith EA, Stuhrmann M. 2006. Dyschromatosis universalis hereditaria as an autosomal recessive disease in five members of one family. Journal of the European Academy of Dermatology and Venereology : JEADV. 20(5):628–629.

Burgess JT, Bolderson E, Adams MN, Baird AM, Zhang SD, Gately KA, Umezawa K, O’Byrne KJ, Richard DJ. 2016. Activation and cleavage of sash1 by caspase-3 mediates an apoptotic response. Cell death & disease. 7(11):e2469.

Burgess JT, Bolderson E, Adams MN, Duijf PHG, Zhang SD, Gray SG, Wright G, Richard DJ, O’Byrne KJ. 2020. Sash1 is a prognostic indicator and potential therapeutic target in non-small cell lung cancer. Scientific reports. 10(1):18605.

Cao L, Zhang R, Yong L, Chen S, Zhang H, Chen W, Xu Q, Ge H, Mao Y, Zhen Q et al. 2021. Novel missense mutation of sash1 in a chinese family with dyschromatosis universalis hereditaria. BMC medical genomics. 14(1):168.

Chen EG, Chen Y, Dong LL, Zhang JS. 2012. Effects of sash1 on lung cancer cell proliferation, apoptosis, and invasion in vitro. Tumour biology : the journal of the International Society for Oncodevelopmental Biology and Medicine. 33(5):1393–1401.

Cui H, Guo S, He H, Guo H, Zhang Y, Wang B. 2020. Sash1 promotes melanin synthesis and migration via suppression of tgf-β1 secretion in melanocytes resulting in pathologic hyperpigmentation. International journal of biological sciences. 16(7):1264–1273.

Derynck R, Turley SJ, Akhurst RJ. 2021. Tgfβ biology in cancer progression and immunotherapy. Nature reviews Clinical oncology. 18(1):9–34.

Goldstein NB, Koster MI, Jones KL, Gao B, Hoaglin LG, Robinson SE, Wright MJ, Birlea SI, Luman A, Lambert KA et al. 2018. Repigmentation of human vitiligo skin by nbuvb is controlled by transcription of gli1 and activation of the β-catenin pathway in the hair follicle bulge stem cells. The Journal of investigative dermatology. 138(3):657–668.

Gong X, Wu J, Wu J, Liu J, Gu H, Shen H. 2017. Correlation of sash1 expression and ultrasonographic features in breast cancer. OncoTargets and therapy. 10:271–276.

Gumienny T, Padgett RW. 2002. The other side of tgf-beta superfamily signal regulation: Thinking outside the cell. Trends in endocrinology and metabolism: TEM. 13(7):295–299.

Gupta A, Sharma Y, Dash KN, Verma S, Natarajan VT, Singh A. 2015. Ultrastructural investigations in an autosomal recessively inherited case of dyschromatosis universalis hereditaria. Acta dermato-venereologica. 95(6):738–740.

Kim DS, Park SH, Park KC. 2004. Transforming growth factor-beta1 decreases melanin synthesis via delayed extracellular signal-regulated kinase activation. The international journal of biochemistry & cell biology. 36(8):1482–1491.

Lin S, Zhang J, Xu J, Wang H, Sang Q, Xing Q, He L. 2012. Effects of sash1 on melanoma cell proliferation and apoptosis in vitro. Molecular medicine reports. 6(6):1243–1248.

Liu JW, Habulieti X, Wang RR, Ma DL, Zhang X. 2021. Two novel sash1 mutations in chinese families with dyschromatosis universalis hereditaria. Journal of clinical laboratory analysis. 35(6):e23803.

Lu C, Liu J, Liu F, Liu Y, Ma D, Zhang X. 2014. Novel missense mutations of abcb6 in two chinese families with dyschromatosis universalis hereditaria. Journal of dermatological science. 76(3):255–258.

Lutz M, Knaus P. 2002. Integration of the tgf-beta pathway into the cellular signalling network. Cellular signalling. 14(12):977–988.

Murphy-Ullrich JE, Suto MJ. 2018. Thrombospondin-1 regulation of latent tgf-β activation: A therapeutic target for fibrotic disease. Matrix biology : journal of the International Society for Matrix Biology. 68-69:28–43.

Ouyang F, Liu X, Liu G, Qiu H, He Y, Hu H, Jiang P. 2020. Long non-coding rna rnf7 promotes the cardiac fibrosis in rat model via mir-543/thbs1 axis and tgfβ1 activation. Aging. 12(1):996–1010.

Pal SK, Nguyen CT, Morita KI, Miki Y, Kayamori K, Yamaguchi A, Sakamoto K. 2016. Thbs1 is induced by tgfb1 in the cancer stroma and promotes invasion of oral squamous cell carcinoma. Journal of oral pathology & medicine : official publication of the International Association of Oral Pathologists and the American Academy of Oral Pathology. 45(10):730–739.

Sethuraman G, Srinivas CR, D’Souza M, Thappa DM, Smiles L. 2002. Dyschromatosis universalis hereditaria. Clinical and experimental dermatology. 27(6):477–479.

Syed V. 2016. Tgf-β signaling in cancer. Journal of cellular biochemistry. 117(6):1279–1287.

Wu N, Tang L, Li X, Dai Y, Zheng X, Gao M, Wang P. 2020. Identification of a novel mutation in sash1 gene in a chinese family with dyschromatosis universalis hereditaria and genotype-phenotype correlation analysis. Frontiers in genetics. 11:841.

Xie J, Zhang W, Zhang J, Lv QY, Luan YF. 2017. Downregulation of sash1 correlates with poor prognosis in cervical cancer. European review for medical and pharmacological sciences. 21(17):3781–3786.

Xing X, Xu Z, Chen L, Jin S, Zhang C, Xiang L. 2022. Tranexamic acid inhibits melanogenesis partially via stimulation of tgf-β1 expression in human epidermal keratinocytes. Experimental dermatology. 31(4):633–640.

Zhang J, Li M, Yao Z. 2017. Updated review of genetic reticulate pigmentary disorders. The British journal of dermatology. 177(4):945–959.

Zhong W, Pan Y, Shao Y, Yang Y, Yu B, Lin Z. 2019. Atypical presentation of dyschromatosis universalis hereditaria with a novel abcb6 mutation. Clinical and experimental dermatology. 44(3):e58–e60.

Zhou D, Wei Z, Deng S, Wang T, Zai M, Wang H, Guo L, Zhang J, Zhong H, He L et al. 2013. Sash1 regulates melanocyte transepithelial migration through a novel gαs-sash1-iqgap1-e-cadherin dependent pathway. Cellular signalling. 25(6):1526–1538.

Zhu Y, Ma Y, Peng H, Gong L, Xiao M, Xiang L, He D, Cao K. 2019. Mir-130b promotes the progression of oesophageal squamous cell carcinoma by targeting sash1. Journal of cellular and molecular medicine. 23(1):93–103.

